# Multicellularity Makes Somatic Differentiation Evolutionarily Stable

**DOI:** 10.1101/010728

**Authors:** Mary E. Wahl, Andrew W. Murray

## Abstract

Many multicellular organisms produce two cell lineages: germ cells, whose descendants form the next generation, and somatic cells which support, protect, and disperse the germ cells. This distinction has evolved independently in dozens of multicellular taxa but is absent in unicellular species. We propose that unicellular, soma-producing populations are intrinsically susceptible to invasion by non-differentiating mutants which ultimately eradicate the differentiating lineage. We argue that multicellularity can prevent the victory of such mutants. To test this hypothesis, we engineer strains of the budding yeast *Saccharomyces cerevisiae* that differ only in the presence or absence of multicellularity and somatic differentiation, permitting direct comparisons between organisms with different lifestyles. We find that non-differentiating mutants overtake unicellular populations but are outcompeted by multicellular differentiating strains, suggesting that multicellularity confers evolutionary stability to somatic differentiation.

**One Sentence Summary:** Using a synthetic biological approach, we show that multicellularity protects species that produce somatic cells from exploitation by common mutants.

## Main Text

Somatic differentiation, a permanent change in gene expression inherited by all of a cell’s descendants, produces somatic cells from a totipotent germ line. Though somatic cells may divide indefinitely, they cannot beget the complete organism and are thus considered non-reproductive. Generation of such sterile cells has clear fitness costs that must be offset by somatic functions which improve the viability or fecundity of germ cells. The absence of a soma in unicellular species (*1*), as well as the persistence of undifferentiated multicellular groups among the volvocine algae (*2*) and cyanobacteria (*3*), has fueled speculation that multicellularity must arise before somatic differentiation can evolve (*4-7*). It has been argued that somatic differentiation is not observed in unicellular species because the potential fitness benefits are insufficient (*6-8*): while soma can contribute motility and protective structures to multicellular organisms, somatic cells in a unicellular species could only benefit the germ line through secretion of useful products into a shared extracellular milieu. However, nutrient exchange between members of microbial consortia (*9, 10*) demonstrates the potential for productive interactions between cell types in the absence of physical adhesion. Benefits associated with somatic differentiation, including reproductive division of labor (*11*) and suppression of germ line mutations through lineage sequestration (*12*), are thus likely accessible to unicellular species.

We propose the alternative hypothesis that unicellular somatic differentiation could offer fitness benefits in a population of genetically identical cells, but remains rare because it is not an evolutionarily stable strategy (*13*). Commonly-occurring mutants that do not differentiate (“cheats”) could take advantage of somatic cell products in the shared media without paying the reproductive costs of differentiation, thus increasing in frequency until their genotype prevails. We further posit that if multicellularity results from cells of a single lineage failing to disperse (rather than cells aggregating from different lineages), differentiating populations may outcompete cheats: although cheats initially arise through mutation in a group with somatic cells, their descendants will eventually be confined to their own multicellular groups composed entirely of cheats and thus cannot benefit from the local accumulation of somatic cell products (*14*).

To test this hypothesis, we designed strains of the budding yeast *Saccharomyces cerevisiae* that produce soma, are multicellular, or both: one strain is a multicellular, differentiating organism and the other two represent both possible intermediates in its evolution from a non-differentiating, unicellular ancestor (Fig. 1A). Employing synthetic strains which differ from one another at only a few, well-defined loci ensures that no undesired variables confound the direct comparison of fitness and evolutionary stability, and permits the study of a unicellular, soma-producing lifestyle not found in nature. Performing experimental tests with living organisms also avoids the potential pitfall of biologically-unrealistic parameter regimes in purely analytical models.

**Fig. 1.**
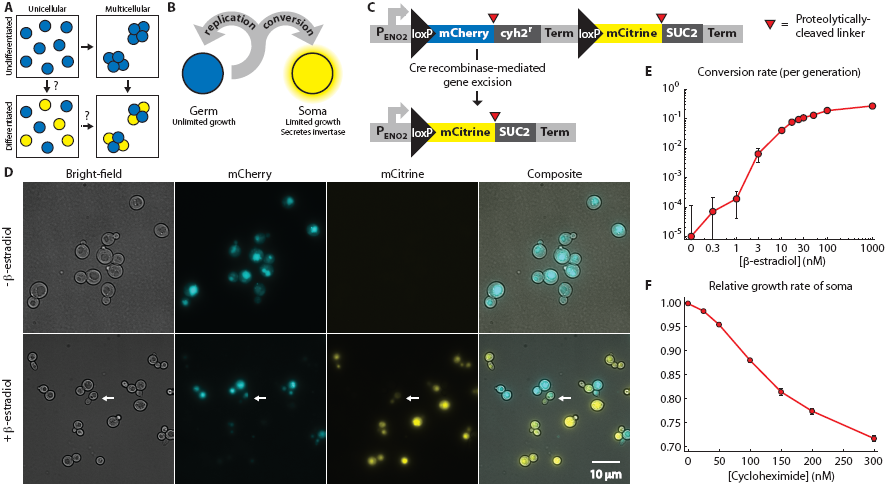
Engineering yeast as a model for somatic differentiation. **(A)** Alternative evolutionary trajectories for the evolution of development. Multicellularity and somatic differentiation have separate biological bases and thus probably evolved sequentially. In some clades, the persistence of likely evolutionary intermediates suggests that multicellularity arose first (solid arrows); no examples of evolution through a unicellular differentiating intermediate (dashed arrows) are known. Somatic cells are colored yellow. **(B)** Schematic of germ-soma division of labor model engineered in our differentiating strains. **(C)** Cell type-defining locus in the differentiating strains. Each pair of cell type-specific proteins (mCherry and Cyh2^r^ for germ cells, mCitrine and Suc2 for somatic cells) is initially expressed as a single polypeptide with a ubiquitin linker (red triangles). Cellular deubiquitinating enzymes cleave this linker post-translationally at its C-terminus (*29*), allowing the two resulting peptides to localize and function independently: for example, Suc2 enters the secretory pathway while mCitrine remains in the cytoplasm. A transcriptional terminator (Term) blocks expression of the somatic cell proteins in germ cells. Cre recombinase-mediated gene excision between loxP sites removes the terminator as well as the germ cell-specific genes. **(D)** The unicellular differentiating strain (yMEW192) during growth in yeast extract-peptone-dextrose media (YPD) before (top) or five hours after addition of 1 μM β-estradiol (bottom). The white arrow indicates a budded cell that has recently undergone conversion and contains both mCherry and mCitrine proteins, and thus fluoresces in both channels (cf. Fig. S1). **(E)** Conversion rates estimated by flow cytometry during growth in YPD containing β-estradiol. Error bars represent 95% confidence intervals of the mean, determined using data obtained from three biological replicates. **(F)** The growth disadvantage of somatic cells relative to germ cells was measured by growing them in co-culture in YPD + cycloheximide and determining the change in the ratio between cell types over multiple rounds of cell division. Error bars represent 95% confidence intervals of the mean, determined using data obtained from three biological replicates.

We mimicked somatic differentiation by engineering fast-growing “germ” cells which can give rise to slower dividing, differentiated “somatic” cells that secrete invertase (Suc2), an enzyme that digests sucrose (which this yeast strain cannot take up directly) into the monosaccharides glucose and fructose, which any cell in the shared medium can then import (*15, 16*) (Fig. 1B). These somatic cells thus perform a digestive function, ensuring the availability of monosaccharides which serve as the sole carbon source during growth in sucrose minimal media. In nature, differentiation is often achieved through multiply-redundant gene regulatory networks that stabilize cell fate (*17*): to simplify our system, we instead made differentiation permanent and heritable by forcing the expression of somatic cell-specific genes to depend on a site-specific recombinase that excises the genes needed for rapid cell proliferation (Fig. 1C).

Germ and somatic cells must be present at a suitable ratio for fast growth on sucrose: germ cells have the higher maximum growth rate, but monosaccharides become limiting when somatic cells are rare. We predicted that the ratio between cell types would reach a steady-state value reflecting the balance between unidirectional conversion of germ cells into somatic cells and the restricted division of somatic cells. We designed tunable differentiation and division rates to allow us to regulate the ratio between cell types and thus control the growth rate of the culture as a whole.

Both features depend on a single, genetically-engineered locus (Fig. 1C). In germ cells, this locus expresses the fluorescent protein mCherry and a gene that accelerates cell division, the cycloheximide resistant (*cyh2^r^*) allele of the ribosomal protein L28 (*ref. 18*); in somatic cells, the locus expresses a different fluorescent protein (mCitrine) and the invertase Suc2. The germ line form is converted to the somatic form by a version of Cre recombinase engineered by Lindstrom et al. (*19*) to be active only in the presence of β-estradiol. Adding β-estradiol to a growing culture induced conversion of germ to somatic cells, apparent as the onset of mCitrine expression and slow loss of mCherry fluorescence by dilution (Figs. 1D and S1; Movie S1). Conversion rates ranged from undetectable levels (< 10^-3^ conversions per cell per generation) to approximately 0.3 conversions per cell per generation as the β-estradiol concentration increased (Figs. 1E and S2A). Expression of the codominant, cycloheximide-sensitive wild-type allele of *CYH2* from its native locus permitted continued growth following *cyh2^r^* excision, but at a reduced rate which depended on cycloheximide concentration. The growth rate deficit of somatic cells ranged from undetectable (< 1%) to nearly 30% as the cycloheximide concentration increased (Figs. 1F and S2B).

The combination of irreversible differentiation and restricted somatic cell division caused cultures to approach a steady-state ratio between the two cell types over time (Fig. 2A). The steady-state fraction of somatic cells increased with the conversion rate and decreased with the somatic cells’ growth disadvantage, as expected (Fig. 2B). The steady-state ratio between cell types could be tuned over four orders of magnitude by altering the cycloheximide and β-estradiol concentrations (Fig. 2B).

**Fig. 2.**
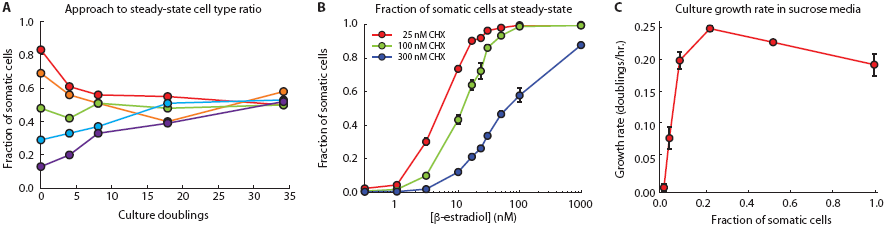
Stable maintenance of both cell types facilitates growth in sucrose + cycloheximide media. **(A)** Representative timecourses of the fraction of somatic cells in cultures of the unicellular differentiating strain (yMEW192 and yMEW192 convertant) initiated at various cell type ratios and passaged in YPD containing 10 nM β-estradiol and 600 nM cycloheximide. **(B)** The steady-state fraction of somatic cells was determined as in (A) for various cycloheximide and β-estradiol concentrations. Error bars represent two standard deviations, calculated using data obtained from three biological replicates. **(C)** Dependence of growth rate on fraction of somatic cells in cultures of the unicellular differentiating strain growing in 0.5% sucrose minimal media. Error bars represent 95% confidence intervals of the mean, determined using data obtained from three biological replicates.

To investigate the ability of invertase secretion from somatic cells to support germ cell proliferation, we determined how the culture’s growth rate on sucrose depended on the ratio between cell types. In the presence of cycloheximide, cultures containing both cell types at an intermediate ratio grew more quickly in sucrose media than cultures of either cell type alone (Fig. 2C), confirming that somatic cells benefit their germ line through invertase secretion.

Clonal multicellularity arises through the maintenance of contact between daughter cells following cytokinesis (*20*). In budding yeast, clonal multicellularity can be produced, either through engineering (*21*) or evolution (*22, 23*), by mutations that prevent degradation of the septum, the specialized part of the cell wall that connects mother and daughter cells after their cytoplasms have been separated by cytokinesis (*24*). Deletion of *CTS1*, a chitinase gene required for septum degradation, causes formation of “clumps” (groups of daughter cells attached through persistent septa) that typically contain 4-30 cells during growth in well-mixed liquid medium (*ref. 25*, Fig. 3A). In the presence of β-estradiol, differentiating strains that lack *CTS1* (*Δcts1*) produced clumps that frequently contained both germ and somatic cells, as evaluated by fluorescence microscopy (Fig. 3A) and flow cytometry (Figs. 3B and 3C). Combining our gene excision-based differentiation system with *CTS1* deletion thus allowed us to produce strains exhibiting all life strategies needed to compare the evolutionary stability of unicellular and multicellular differentiation.

**Fig. 3.**
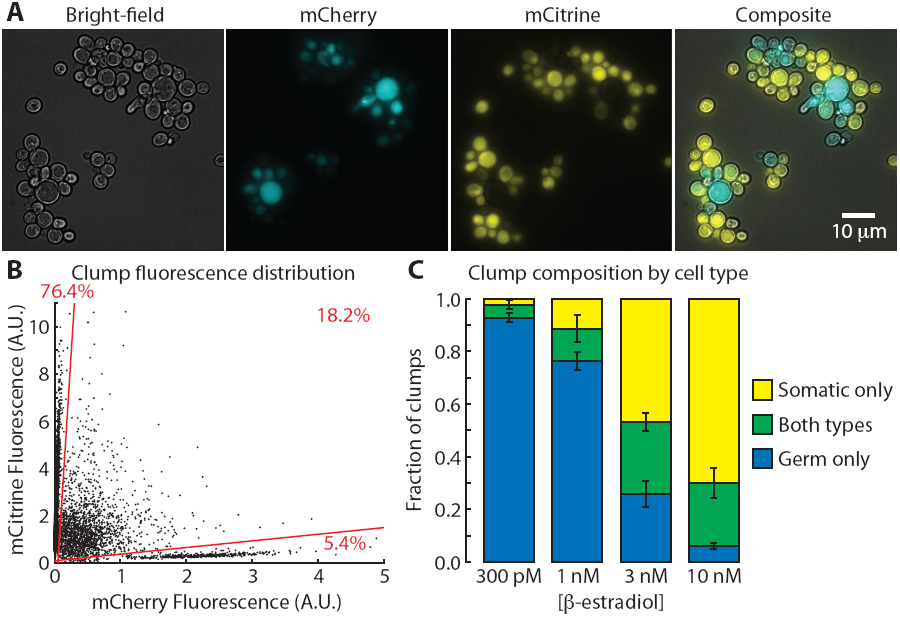
*Δcts1* strains form multicellular clumps containing both cell types. **(A)** Representative image of clump size and cell type composition in a well-mixed liquid culture of the *Δcts1* multicellular differentiating strain (yMEW208) at a steady-state cell type ratio in sucrose minimal media containing 10 nM β-estradiol and 150 nM cycloheximide. **(B)** Representative distribution of clump mCherry and mCitrine fluorescence for the multicellular differentiating strain (yMEW208) at a steady-state cell type ratio in sucrose minimal media containing 10 nM β-estradiol and 150 nM cycloheximide. Gating of clumps by cell type composition (lower sector: germ cells only; middle sector: both cell types; top sector: somatic cells only) is shown in red. **(C)** The fraction of clumps containing one or both cell types was determined as in (B) for cultures of the multicellular differentiating strain (yMEW208) at a steady-state cell type ratio in sucrose minimal media containing 150 nM cycloheximide at the indicated β-estradiol concentrations. Error bars represent 95% confidence intervals of the mean, determined with data obtained from six biological replicates.

Somatic cells in unicellular strains can only benefit germ cells by secreting useful products into a shared medium; non-differentiating cheats (e.g., Cre^-^ germ cells) and germ cells in well-mixed media have equal access to somatic cell products, but cheats do not pay the reproductive toll of differentiation. We therefore predicted that cheats would enjoy a fitness advantage over germ cells, allowing them to invade unicellular, differentiating populations (Fig. 4A, top). In multicellular species, however, significant local accumulation of somatic cell products (in our experiment, monosaccharides) within multicellular groups can give differentiating lineages an advantage over cheats as long as the benefit of better nutrition overcomes the cost of producing slower-replicating, somatic cells (*21*). In clonally multicellular species, such as our *Δcts1* strain, novel cheats arising by mutation will eventually be segregated into cheat-only groups by cell division and group fragmentation. We hypothesized that cheats would then experience reduced access to somatic cell products, potentially negating their growth advantage over germ cells (Fig. 4A, bottom).

**Fig. 4.**
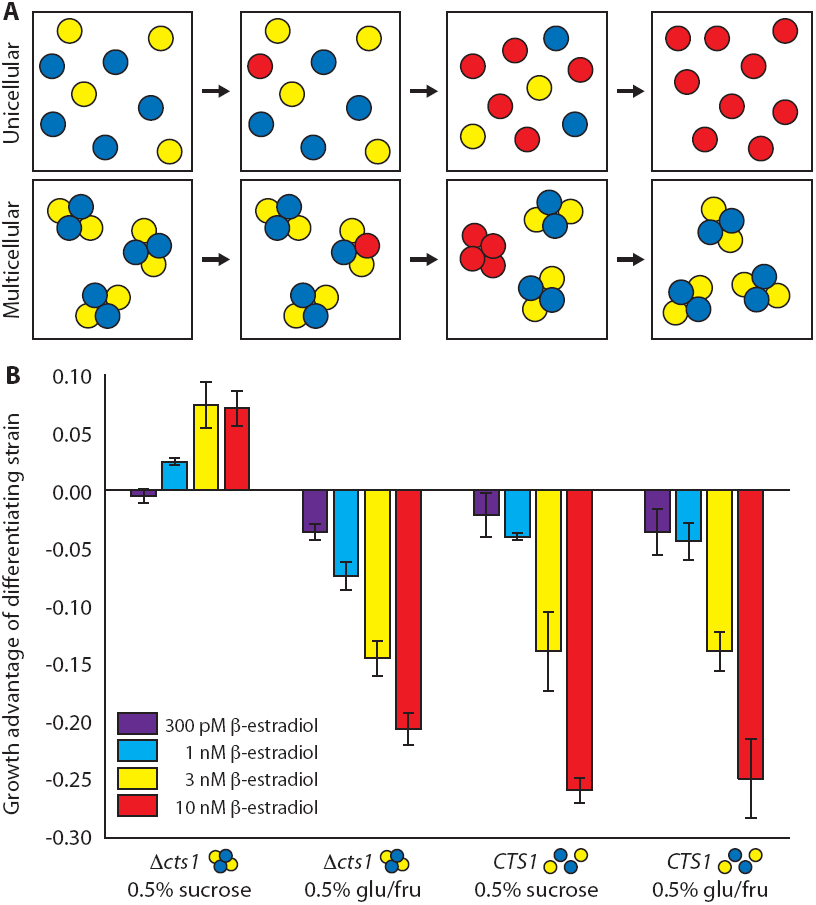
Multicellular differentiating strains resist invasion by Cre^-^ cheats. **(A)** Hypothesized evolutionary outcomes for novel non-differentiating mutants (red) in unicellular (top) and multicellular (bottom) populations. Germ cells are blue and somatic cells are yellow. **(B)** Growth advantages of differentiating strains (unicellular: yMEW192, multicellular: yMEW208) relative to cheaters (unicellular: yMEW193, multicellular: yMEW209) in minimal media containing 150 nM cycloheximide. Error bars represent 95% confidence intervals of mean growth advantages, determined using data obtained from three biological replicates. Growth advantages of the multicellular strain growing in sucrose minimal medium at 1 nM, 3 nM, and 10 nM β-estradiol are significantly greater than zero (p < 10^-3^, one-tailed t-test); growth advantages of the unicellular strain growing in sucrose are significantly less than zero at all β-estradiol concentrations (p < 10^-6^, one-tailed t-test).

To test this prediction, we introduced cheats into unicellular or multicellular differentiating cultures and investigated their fate. We mixed differentiating cultures expressing a third fluorescent protein, Cerulean, with cheats that lack this third color and cannot differentiate because they lack the recombinase whose action gives rise to somatic cells (Cre^-^). We monitored the relative frequency of the two strains over a series of growth and dilution cycles. In sucrose media, cheats invaded unicellular, differentiating populations but were outcompeted in multicellular, differentiating populations (Fig. 4B). The growth advantage of multicellular, differentiating strains was nullified in monosaccharide-containing media, where somatic cells should confer no fitness advantage to clumps (Fig. 4B). Moreover, the growth advantage of differentiating strains depended strongly on the conversion rate (β-estradiol concentration): higher conversion rates were advantageous only in the multicellular differentiating case (Fig. 4B), where they increased the fraction of somatic cells overall as well as the fraction of clumps containing at least one somatic cell (Fig. 3C). In unicellular cultures grown on sucrose, or in cultures grown on glucose (unicellular or multicellular), increasing the conversion rate increased the growth advantage of cheats by reducing the growth rate of the germ cell population (Fig. 4B).

Our results show that unicellular, soma-producing strains are evolutionarily unstable to invasion by non-differentiating cheats. This finding is unlikely to depend on the specific molecular mechanisms that effect differentiation or allow somatic cells to assist germ cells: any form of differentiation in a single-celled species would require that the two cell types exchange resources through a shared medium. From the spontaneous mutation rate in budding yeast [≈ 3 × 10^-10^ per base pair per cell division (*26*)] and the size of the recombinase gene, we estimate mutations that inactivate our engineered recombination system would occur in about one out of every 10^7^ cell divisions; this is likely an underestimate of the frequency of inactivating mutations in natural differentiation, which typically requires more loci and thus presents a larger target for mutation (*17*). Because this frequency is high relative to typical microbial population sizes, cheats would thus likely arise and sweep to fixation shortly after the appearance of unicellular somatic differentiation, explaining the absence of extant species with this life strategy. The short persistence time of unicellular differentiating species makes them an unlikely intermediate in the evolution of development relative to undifferentiated multicellular species, which have persisted for hundreds of millions to billions of years in some clades (*1-3*). We cannot, however, rule out the possibility that the transient existence of a unicellular differentiating species might suffice for the secondary evolution of multicellularity, which has been observed experimentally in small populations on the timescale of weeks (*22, 27*). In any case, the differentiating phenotype cannot be stably maintained against cheats until clonal multicellularity evolves.

Our study demonstrates that synthetic biology can directly test hypotheses about evolutionary transitions, complementing retrospective inference through comparison of existing species, experimental evolution, mathematical modeling, and simulation. We note that other major evolutionary transitions, including the appearance of body plans (spatially-ordered arrangements of cell types) and life cycles (temporal sequences of growth and dispersion), could be studied through experimental evolution or further engineering of the strains described above. Furthermore, our differentiating strain provides an experimentally-tractable version of a common simplifying assumption in population genetics: independent deleterious mutations are often modeled as having the same fitness cost (*28*). In our strains, *cyh2^r^* excision is a form of irreversible mutation that always produces the same fitness disadvantage, but the magnitude of the fitness cost can be experimentally varied by altering the cycloheximide concentration. Thus engineered organisms, including those we have developed, can permit robust experimental testing of a wide variety of outstanding hypotheses in evolutionary biology.

## Acknowledgments

We thank Derek Lindstrom and Dan Gottschling for providing the estradiol-inducible Cre construct *P*_*SCW11*_-*cre-EBD78*; Alex Schier, Michael Desai, Michael Laub, Cassandra Extavour, David Haig, and members of the Murray and Nelson labs for helpful discussions during manuscript preparation; and Beverly Neugeboren, Linda Kefalas, and Sara Amaral for research support. This work was supported in part by National Science Foundation and Department of Defense National Defense Science and Engineering Graduate Research Fellowships and by NIH/NIGMS grant GM068763.

## Supplementary Materials

### Materials and Methods

#### Plasmid construction

A complete list of plasmids used in this study is given in Table S1; plasmid maps and sequences are available upon request. Constructs were cloned using standard methods (*30, 31*) and common base vectors (*32, 33*). Yeast-optimized sequences encoding the fluorescent proteins Cerulean (*34*), mCitrine (*35*) and mCherry (*36*) (here denoted yCerulean, ymCitrine, and ymCherry) were generously provided by Nicholas Ingolia.

*loxP*, the binding site of Cre recombinase, is 34 bp long and contains inverted repeats that can interfere with PCR-based cloning strategies. We reasoned that inclusion of this inverted repeat within a transcript could also form a hairpin or otherwise interfere with translation. We therefore introduced the *loxP* sequence into a yeast artificial intron (*38*) to facilitate cloning. The *loxP*-containing artificial intron (*AI*) was then added to the 5’ end of the *ymCherry* open reading frame and the *ADH1* terminator by fusion PCR and cloned into the base vector pFA6a-HIS3MX6 to generate pMEW56. The constitutive actin promoter *P*_*ACT1*_ was then introduced into pMEW56 by ligation-independent cloning, producing pMEW61.

A truncated version of the artificial intron which lacked the 5’ splice site was added to the 5’ end of the *SUC2* open reading frame and terminator by fusion PCR and cloned into pRS402 to generate pMEW54. The *AI-SUC2-T_SUC2_* amplicon was then cloned into pMEW61 by restriction digestion and ligation to produce pMEW63.

Constructs introduced by conventional plasmid integration are subject to spontaneous loop-out or increase in copy number; our application required the introduction of exactly one copy of the construct, motivating the development of an integration strategy based on gene replacement (*39*). Both constructs of interest (Figs. 1C and S3A) contained sequences homologous to the *SUC2* open reading frame and terminator at their 3’ ends. We therefore designed a base vector containing sequence homologous to the 5’ end of the *SUC2* locus so that appropriate restriction digestions would release a fragment that would interact at both ends of the *SUC2* locus, replacing the *SUC2* gene with the engineered DNA. Three hundred base pairs of the *SUC2* promoter region were therefore introduced into the base vector pRS402 5’ to its *ADE2* marker by ligation-independent cloning to create pMEW71. The *P*_*ACT1*_-*AI-ymCherry-T*_*ADH1*_-*AI-SUC2-T*_*SUC2*_ fragment of pMEW63 was then cloned into pMEW71 3’ to the *ADE2* marker such that a Bsu36I/BamHI digestion fragment of the resulting plasmid (pMEW73) would contain the *ADE2* marker, the construct of interest, and homology to the *SUC2* locus at either end to direct homologous recombination.

An intron-free version of the ribosomal protein L28-encoding gene *CYH2* containing a previously-described (*18*) mutation conferring cycloheximide resistance, *cyh2-N38K* (*cyh2^r^*), was produced by fusion and site-specific, mutagenic PCR. This construct was then introduced in place of *ymCherry-T_ADH1_* in pMEW61 to generate pMEW67. To strengthen the cycloheximide resistance conferred by this codominant allele (*40*), the actin promoter was replaced with the stronger constitute enolase promoter *P*_*ENO2*_ (*ref. 41*) by ligation-independent cloning, generating pMEW76.

To express a fluorescent marker and a second protein from a single open reading frame with minimal disruption of localization or function, a ubiquitin moiety (*UBQ*) linker was included in both gene fusions (*ymCherry-UBQ-cyh2^r^* and *ymCitrine-UBQ-SUC2*). Cleavage at the C-terminus of this linker by native ubiquitin-specific proteases (*29*) after translation produces two peptides: a fluorescent protein with C-terminal ubiquitin moiety (e.g., ymCherry-Ubq and ymCitrine-Ubq) which does not significantly impact its stability, and the native form of the second protein (e.g., Cyh2^r^ and Suc2 with no N-terminal tags). The first ubiquitin moiety of*UBI4* was amplified and fused to the 3’ end of the *AI-mCherry* construct, then introduced into pMEW76 by ligation-independent cloning to generate pMEW81. *P*_*ENO2*_ was then added to the 5’ end of the *AI-ymCherry-UBQ-cyh2^r^-T_CYH2_* construct by fusion PCR, and the resulting amplicon was introduced into pMEW73 by restriction digest to create pMEW82.

An *AI-ymCitrine-UBQ-SUC2-T_SUC2_* construct was generated by replacing the *ymCherry* open reading frame in pMEW81 with *ymCitrine* by ligation-independent cloning (producing the intermediate plasmid pMEW83), then replacing *cyh2^r^-T_CYH2_* with *SUC2-T_SUC2_* by ligation-independent cloning to generate pMEW84. The *AI-ymCitrine-UBQ-SUC2-T_SUC2_* fragment was then introduced into pMEW82 by restriction digest to create pMEW90, which contains the complete construct used to produce the *cyh2^r^* excision strain.

#### Strain construction

A complete list of strains used in this study is given in Table S2. All constructs and markers were introduced via homologous recombination using a standard polyethylene glycol, lithium acetate, and Tris-EDTA yeast transformation protocol (*30*); integration at the desired loci was confirmed by diagnostic PCR. All strains are constructed in a W303 background in which a loss-of-function mutation in *BUD4* has been corrected as previously described (*21, 42*).

The β-estradiol-inducible Cre construct previously described by Lindstrom et al. (*19*) was introduced by homologous recombination at the *HO* locus. The P_ACT1_-yCerulean construct was amplified by PCR and integrated at the *LEU2* locus. The insert of pMEW90 was released by Bsu36I/BamHI digestion prior to transformation. The *CTS1* open reading frame was deleted by adding 40 bp of *CTS1* locus homology to each end of the *HIS3MX6* marker [containing the open reading frame of *Schizosaccharomyces pombe* gene *HIS5*, which complements *S. cerevisiae his3* mutations (*43*)] by PCR amplification prior to transformation. yMEW192 convertants were isolated by diluting and plating cultures of yMEW192 which had previously been grown in media containing β-estradiol to induce conversion.

#### Media and growth conditions

Minimal media and yeast extract-peptone-dextrose (YPD) media were produced as described elsewhere (*21*). β-estradiol (Sigma-Aldrich E8875) and cycloheximide (Sigma Aldrich C7698) were resuspended at 1 mM in ethanol and stored at −80°C for less than one year; media containing these reagents were stored less than one week at room temperature. Shaken liquid cultures and agar plates were grown at 30°C. Sucrose was dissolved in solution without heating and stored at 4°C less than one month to limit spontaneous hydrolysis.

#### Fluorescence microscopy

Still images and time-lapse videos were collected at room temperature using an Eclipse Ti-E inverted microscope (Nikon Instruments, Melville, NY) with a 60x Plan Apo VC 1.4NA oil objective, a Photometrics CoolSNAP HQ camera (Roper Scientific), and appropriate filters for yeast-optimized Cerulean (ex=436/20 nm, em=480/40 nm), mCitrine (ex=500/20 nm, em=535/30 nm), and mCherry (ex=562/40 nm, em=641/75 nm) fluorescent proteins. The image processing program Fiji (*44*) was used to produce composite images and generate video files.

Time-lapse videos of monolayer growth were collected using a CellASICs Y04C ONIX Live Cell Imaging microfluidics flow chamber (EMD Millipore, Billerica, MA) pre-treated by perfusion of concanavalin A solution (1 mg/mL concanavalin A, 1 mM CaCl_2_, 10 mM NaHPO_4_ at pH 6) for 5 minutes at 2 psi, followed by washout with growth media for 5 minutes at 2 psi. Cells of a late log-phase culture (approx. 5 × 10^6^ cells/mL) were loaded into the chamber at 5 psi for 10 seconds. Metamorph 7.7 (Molecular Devices, Sunnyvale, CA) with Nikon Perfect Focus System (Nikon Instruments, Melville, NY) was used to acquire images at multiple stage positions at exposure times. For videos, fluorescence and bright-field images were collected at multiple stage positions at fifteen-minute intervals.

#### Flow cytometry

An LSR Fortessa (BD Biosciences, San Jose, CA) with appropriate filters for yeast-optimized Cerulean (ex=440 nm, em=470/20 nm), mCitrine (ex=488 nm, em=530/30 nm), and mCherry (ex=561 nm, em=610/20 nm) was used to perform flow cytometry on well-vortexed samples of culture aliquots. For unicellular culture samples, appropriate forward and side scatter gating were used to eliminate events corresponding to multiple cells.

#### Conversion timecourses and conversion rate assays

Conversion rate timecourses were performed by adding β-estradiol at the indicated concentration to cultures of pure germ cells (yMEW192) which were pre-grown in log phase in YPD. Cultures were maintained in log phase for additional growth while aliquots were collected for analysis by flow cytometry. At the indicated time point, cultures were washed twice with phosphate-buffered saline to remove β-estradiol and resuspended in YPD for additional growth before a final flow cytometry analysis.

For conversion rate determinations, β-estradiol was added to pure cultures of germ cells (yMEW192) pre-grown in log phase in YPD (Fig. S2A). Pure cultures of germ cells were maintained in parallel to permit calculation of the number of germ cell generations elapsed through cell density measurement with a Coulter counter (Beckman Coulter, Danvers, MA). Aliquots collected at indicated time points were washed twice in phosphate-buffered saline solution to remove β-estradiol, then resuspended in YPD for continued growth, allowing recently-converted somatic cells to dilute out remaining mCherry so that cell types could be unambiguously distinguished by flow cytometry. Linear regression of the log fraction of cells which were germ cells vs. the number of germ cell generations elapsed was used to determine a 95% confidence interval of the conversion rate for each experiment using the fit() and confint() functions in MATLAB (Mathworks, Natick, MA). The conversion rates reported in Figure 1E correspond to the weighted arithmetic mean and corresponding variance calculated with data obtained from three or more independent experiments.

#### Fitness assays for relative growth difference between cell types

The growth rate difference between cell types was determined using a fitness assay protocol described previously (*45*). Briefly, pure cultures of germ (yMEW192) and somatic (yMEW192 convertant) cells pre-grown in log phase in YPD containing cycloheximide were combined and maintained in log phase through multiple growth and dilution cycles in like media (Fig. S2B). Aliquots taken at each dilution step were used to determine the fraction of each cell type by flow cytometry. Pure cultures of germ (yMEW192) cells were maintained in log phase in parallel to determine the number of germ cell generations elapsed through cell density measurements with a Beckman Coulter counter. The 95% confidence interval for the slope of the linear regression line of log(somatic cells/germ cells) vs. the number of germ cell generations elapsed was determined as described above. The growth rate differences reported in Figure 1F represent the weighted arithmetic means and corresponding variances of the slopes calculated in three independent experiments.

#### Steady-state ratio assays

Pure cultures of germ cells (yMEW192) and somatic cells (yMEW192) were mixed to initiate cultures from a range of cell type ratios. Cultures were washed with phosphate-buffered saline and resuspended in YPD containing β-estradiol and cycloheximide at the indicated concentrations. Aliquots were taken at the indicated time points to determine the number of culture doublings since initiation (determined from Coulter counter measurements of culture density) and the fraction of somatic (mCitrine^+^ mCherry^-^) cells in the culture. Cultures typically converged on a steady-state ratio after 30-40 culture doublings.

#### Growth rate assays in sucrose minimal media

Cultures of converting strains at their steady-state ratios were washed twice with phosphate-buffered saline and resuspended at approximately 10^5^ cells/mL in minimal media containing 0.5% sucrose and the concentrations of cycloheximide and β-estradiol in which they had been growing previously. Pure cultures of the Cre^-^ strain yMEW193 and the yMEW192 convertant were also used to represent pure cultures of germ and somatic cells of the differentiating strain, respectively. After an acclimation period of approximately ten hours, culture density measurements were taken regularly with a Coulter counter and a 95% confidence interval of the slope determined by linear regression of log(culture density) vs. time. The reported growth rates (Fig. 2C) were obtained from the weighted arithmetic means of slopes and their corresponding variances estimated from three or more independent experiments.

#### Determination of the distribution of cell type composition in clumps

Cultures of *Δcts1* strains growing in minimal media containing 0.5% sucrose and the indicated concentrations of cycloheximide and β-estradiol were analyzed by flow cytometry. Clumps in these cultures typically ranged in size from 4-30 clumps; forward and side scatter gating could therefore not be used to eliminate events consisting of two or more clumps. Instead, cultures were diluted and vortexed thoroughly prior to flow cytometry to reduce the probability of multiple clumps being counted as a single event. The efficacy of this strategy was checked by combining pure cultures of germ and somatic cells in the absence of β-estradiol: the fraction of mCherry+ mCitrine+ events in the mixed culture was less than 1% (data not shown). Clumps containing only germ cells had negligible fluorescence in the mCitrine channel and thus formed a distinguishable population; likewise, clumps containing only somatic cells had negligible fluorescence in the mCherry channel. Clumps with non-negligible fluorescence in both channels were considered to contain both cell types.

#### Competition assays

In competition assays, Cerulean^+^ differentiating strains (unicellular: yMEW192; multicellular: yMEW208) were mixed with Cerulean^-^ Cre^-^ reference strains (unicellular: yMEW193; multicellular: yMEW209) at a 1:1 ratio and passaged through five cycles of growth and dilution in indicated media. Aliquots taken at each dilution step were analyzed by flow cytometry to determine the ratio between Cerulean^+^ and Cerulean^-^ events (cells or clumps). The 95% confidence interval of the slope of the linear regression line of log(Cerulean^+^ events/Cerulean^-^ events) vs. the number of culture doublings elapsed was determined as described above. The growth advantages of the differentiating strain reported in Figure 4B represent the weighted arithmetic means and corresponding variances of the slopes calculated in three or more independent experiments.

**Fig. S1.**
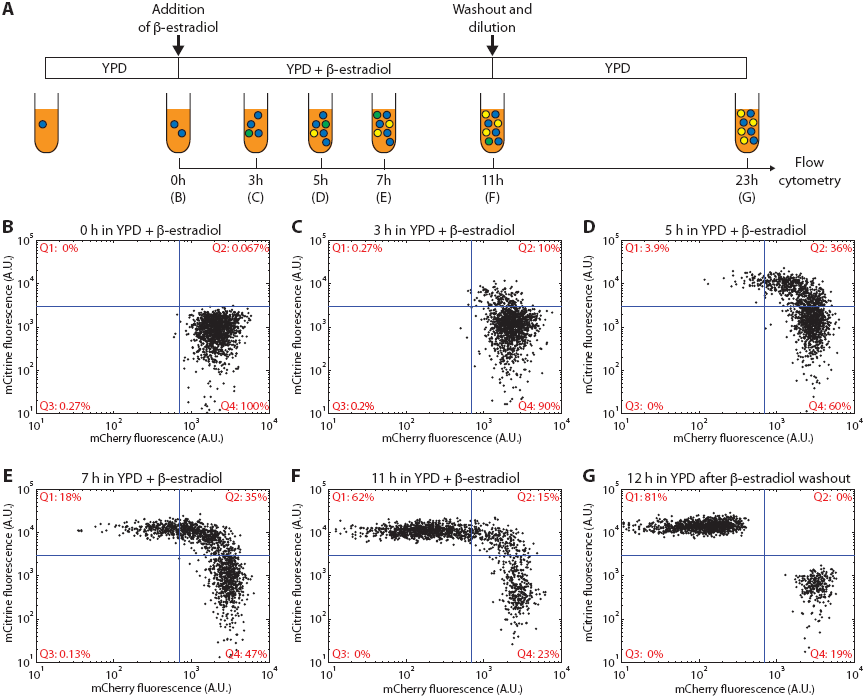
Flow cytometry timecourse of conversion in a *cyh2^r^* excision strain. **(A)** Schematic of the experiment. A culture of germ cells (blue) of the *cyh2r* excision strain (yMEW192) was pre-grown in log phase in yeast extract-peptone-dextrose (YPD) media. Immediately after addition of 1 μM β-estradiol (**B**), and at the indicated timepoints during subsequent growth (**C**-**F**), aliquots were collected for flow cytometry. Converted cells began to express mCitrine and lost mCherry fluorescence gradually by dilution through growth. After 11 hours in 1 μM β-estradiol, cells were washed and transferred to fresh YPD media: twelve hours of further growth and division diluted remaining mCherry from mCitrine^+^ somatic cells, permitting the two cell types to be unambiguously distinguished (**G**). Reversion to the germ cell state, as defined by recovery of mCherry fluorescence, was never observed by flow cytometry, time-lapse microscopy in flow chambers, or fluorescence colony imaging (data not shown).

**Fig. S2.**
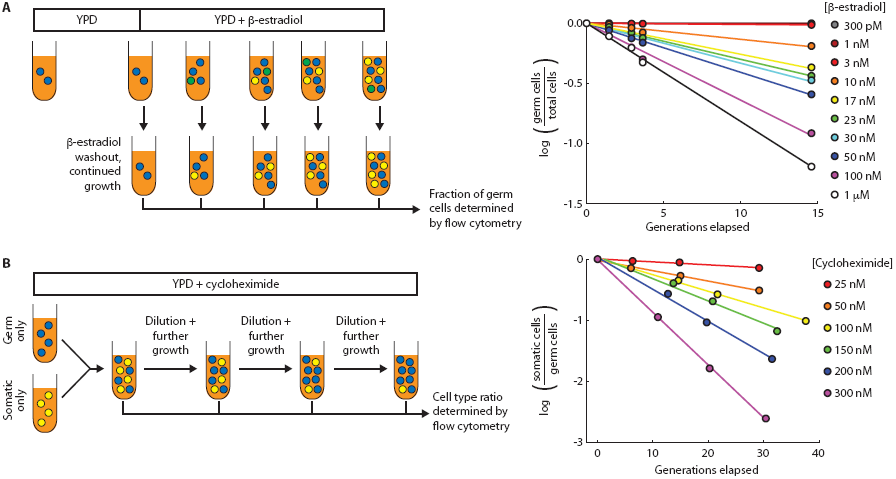
Measurement of conversion and relative growth rates. **(A)** Measurement of conversion rates by flow cytometry. Aliquots were collected at various time points after induction of conversion to determine the number of generations elapsed since β-estradiol addition as well as the fraction of mCherry^+^ mCitrine^-^ (germ) cells within the culture. β-estradiol washout and continued growth permits unambiguous discrimination of germ and somatic cells by flow cytometry (cf. Fig. S1G). Representative data are shown at right. In the absence of cycloheximide, the fraction of germ cells is predicted to fall exponentially according to the conversion rate. Slopes from best-fit linear regression lines were used to calculate conversion rates (Fig. 1E). **(B)** Measurement of relative growth rate differences through competition fitness assays. Pure cultures of germ and somatic cells were combined at a 1:1 ratio and passaged in log phase in YPD + cycloheximide. Aliquots were collected at each dilution to determine the ratio between cell types by flow cytometry. Representative data are shown at right. Slopes calculated from best-fit regression lines were used to determine differences in relative growth rates (Fig. 1F).

**Table S1:** Plasmids used in the construction of transgenic yeast strains.

| Plasmid | Base Vector | Insert | Strains with insert integrated by homologous recombination |
| --- | --- | --- | --- |
| pDL12 | from ref. 19 | *P<sub>SCW11</sub>-cre-EBD78-T<sub>CYC1</sub>* | - |
| pMEW54 | pRS402 | *AI-SUC2-T<sub>SUC2</sub>* | - |
| pMEW56 | pFA6a-HIS3MX6 | *AI-ymCherry-T<sub>ADH1</sub>* | - |
| pMEW61 | pFA6a-HIS3MX6 | *P<sub>ACT1</sub>-AI-ymCherry-T<sub>ADH1</sub>* | - |
| pMEW63 | pFA6a-HIS3MX6 | *P<sub>ACT1</sub>-AI-ymCherry-T<sub>ADH1</sub>-AI-SUC2-T<sub>SUC2</sub>* | - |
| pMEW67 | pFA6a-HIS3MX6 | *P<sub>ACT1</sub>-AI-cyh2'-T<sub>CYH2</sub>* | - |
| pMEW71 | pRS402 | *5'SUC2* | - |
| pMEW73 | pRS402 | *5'SUC2 P<sub>ACT1</sub>-AI-ymCherry-T<sub>ADH1</sub>-AI-SUC2-T<sub>SUC2</sub>* | - |
| pMEW76 | pFA6a-HIS3MX6 | *P<sub>ENO2</sub>-AI-cyh2'-T<sub>CYH2</sub>* | - |
| pMEW81 | pFA6a-HIS3MX6 | *AI-ymCherry-UBQ-cyh2'-T<sub>CYH2</sub>* | - |
| pMEW82 | pRS402 | *5'SUC2 P<sub>ENO2</sub>-AI-ymCherry-UBQ-cyh2'-T<sub>CYH2</sub>-AI-SUC2-T<sub>SUC2</sub>* | - |
| pMEW83 | pFA6a-HIS3MX6 | *AI-ymCitrine-UBQ-cyh2'-T<sub>CYH2</sub>* | - |
| pMEW84 | pFA6a-HIS3MX6 | *AI-ymCitrine-UBQ-SUC2-T<sub>SUC2</sub>* | - |
| pMEW90 | pRS402 | *5'SUC2 P<sub>ENO2</sub>-AI-ymCherry-UBQ-cyh2'-T<sub>CYH2</sub>-AI-ymCitrine-UBQ-SUC2-T<sub>SUC2</sub>* | yMEW192<br>yMEW193<br>yMEW208<br>yMEW209 |

**Table S2:** Transgenic yeast strains used in this study.

| Strain name | Genotype description |
| --- | --- |
| yMEW17 | W303 MATa *BUD4 bar1 leu2-3 can1-100 ura3-1 ade2-1 his3-11 trp1-1* |
| yMEW192 | yMEW17 *TRP1 HIS3* $\Delta ho::P_{SCW11}$-cre-EBD78::NATMX4 $\Delta leu2::HPHMX4::P_{ACT1}$-yCerulean-*T<sub>ADH1</sub>*<br>*ADE2::P<sub>ENO2</sub>*-Al-ymCherry-UBQ-cyh2'-*T<sub>CYH2</sub>*-Al-ymCitrine-UBQ-SUC2-*T<sub>SUC2</sub>* |
| yMEW192<br>convertant | yMEW17 *TRP1 HIS3* $\Delta ho::P_{SCW11}$-cre-EBD78::NATMX4 $\Delta leu2::HPHMX4::P_{ACT1}$-yCerulean-*T<sub>ADH1</sub>*<br>*ADE2::P<sub>ENO2</sub>*-Al-ymCitrine-UBQ-SUC2-*T<sub>SUC2</sub>* |
| yMEW193 | yMEW17 *TRP1 HIS3 ADE2::P<sub>ENO2</sub>*-Al-ymCherry-UBQ-cyh2'-*T<sub>CYH2</sub>*-Al-ymCitrine-UBQ-SUC2-*T<sub>SUC2</sub>* |
| yMEW208 | yMEW17 *TRP1* $\Delta ho::P_{SCW11}$-cre-EBD78::NATMX4 $\Delta leu2::HPHMX4::P_{ACT1}$-yCerulean-*T<sub>ADH1</sub>*<br>*ADE2::P<sub>ENO2</sub>*-Al-ymCherry-UBQ-cyh2'-*T<sub>CYH2</sub>*-Al-ymCitrine-UBQ-SUC2-*T<sub>SUC2</sub>* $\Delta cts1::HIS3$ |
| yMEW209 | yMEW17 *TRP1 ADE2::P<sub>ENO2</sub>*-Al-ymCherry-UBQ-cyh2'-*T<sub>CYH2</sub>*-Al-ymCitrine-UBQ-SUC2-*T<sub>SUC2</sub>* $\Delta cts1::HIS3$ |

**Movie S1:** Time-lapse imaging of *cyh2^r^* excision strain colony formation.

Representative time-lapse imaging of cells of the *cyh2^r^* excision strain (yMEW192) growing in YPD containing 1 μM β-estradiol within a flow chamber. Top left: Bright-field; top right: mCherry; bottom left: mCitrine; bottom right: composite.

